# Sequence-based modelling of bacterial genomes enables accurate antibiotic resistance prediction

**DOI:** 10.1101/2024.01.03.574022

**Authors:** Maciej Wiatrak, Aaron Weimann, Adam Dinan, Maria Brbić, R. Andres Floto

## Abstract

Rapid detection of antibiotic-resistant bacteria and understanding the mechanisms underlying antimicrobial resistance (AMR) are major unsolved problems that pose significant threats to global public health. However, existing methods for predicting antibiotic resistance from genomic sequence data have had limited success due to their inability to model epistatic effects and generalize to novel variants. Here, we present GeneBac, a deep learning method for predicting antibiotic resistance from DNA sequence through the integration of interactions between genes. We apply GeneBac to two distinct bacterial species and show that it can successfully predict the minimum inhibitory concentration (MIC) of multiple antibiotics. We use the WHO Mycobacterium tuberculosis mutation catalogue to demonstrate that GeneBac accurately predicts the effects of different variants, including novel variants that have not been observed during training. GeneBac is a modular framework which can be applied to a number of tasks including gene expression prediction, resistant gene identification and strain clustering. We leverage this modularity to transfer learn from the transcriptomic data to improve performance on the MIC prediction task.

## 1 Main

Antimicrobial Resistance (AMR) is a major global issue, with more than 1.27 million deaths accredited to it annually [1, 2]. A significant obstacle in combatting AMR is a rapid and accurate diagnosis of antibiotic susceptibility or resistance, allowing for the right drug to be administered. Moreover, as microbial agents develop resistance mechanisms against existing antibiotics, it is crucial to understand the molecular mechanisms leading to the resistance.

Whole genome sequence-related diagnostic tests coupled with statistical methods, have proven effective in predicting antibiotic resistance in bacteria [3–8]. However, these approaches exhibit notable limitations. They struggle to consider the combinatorial effects of multiple genetic variants and epistatic effects [9, 10], hindering a comprehensive understanding of resistance mechanisms. Furthermore, their inability to assess the impact of newly observed or rare variants poses a challenge in staying ahead of evolving resistance patterns. Additionally, these diagnostic tests ignore the genetic context of a variant, such as a variant falling in a binding site and may be influenced by population structure, impacting the generalisability of their predictions [11]. Addressing these limitations remains crucial for refining the accuracy and applicability of whole genome sequence-based antibiotic resistance predictions. Recently, machine learning methods have been shown to outperform statistical methods in predicting resistance to multiple drugs on *Mycobacterium tuberculosis* [12–16]. However, these models still fail to account for the interactions between genes and ignore the genetic context of variants. The latter is particularly important to understand the mechanisms leading to resistance and to be able to generalise to variants not observed during training, which is crucial considering the rapid rate of mutation in bacteria. Finally, existing methods treat the antibiotic resistance problem as a classification task, ignoring the fact that the minimum inhibitory concentration (MIC) distributions are usually not binary (Supp. Fig. 1a-b), which may lead to an inappropriate diagnosis in the clinical setting [17, 18].

In this work, we propose GeneBac, a deep learning model for the prediction of minimum inhibitory concentration (MIC) of antibiotics in bacteria. GeneBac is a DNA sequence model [19–23] that accounts for the genetic context of a variant, such as the Transcription Factor (TF)-DNA binding motif by directly leveraging the sequence of nucleotides as input. GeneBac has two main components, a sequence-based model capable of capturing the variation in the DNA sequence of a gene and a graph neural network (GNN), modelling gene-gene interactions in a single strain (Fig. 1a). Through the use of a DNA sequence model, GeneBac is capable of assessing the impact of novel mutations and accounting for the variants’ genetic context. Moreover, by leveraging a GNN model, GeneBac incorporates epistatic effects for antibiotic resistance prediction. Since GeneBac uses DNA with variants directly integrated into the sequence as input, it has a unique ability to generalise to new variants that were not observed during training and shows robustness to the bacterial population structure.

**Fig. 1.**
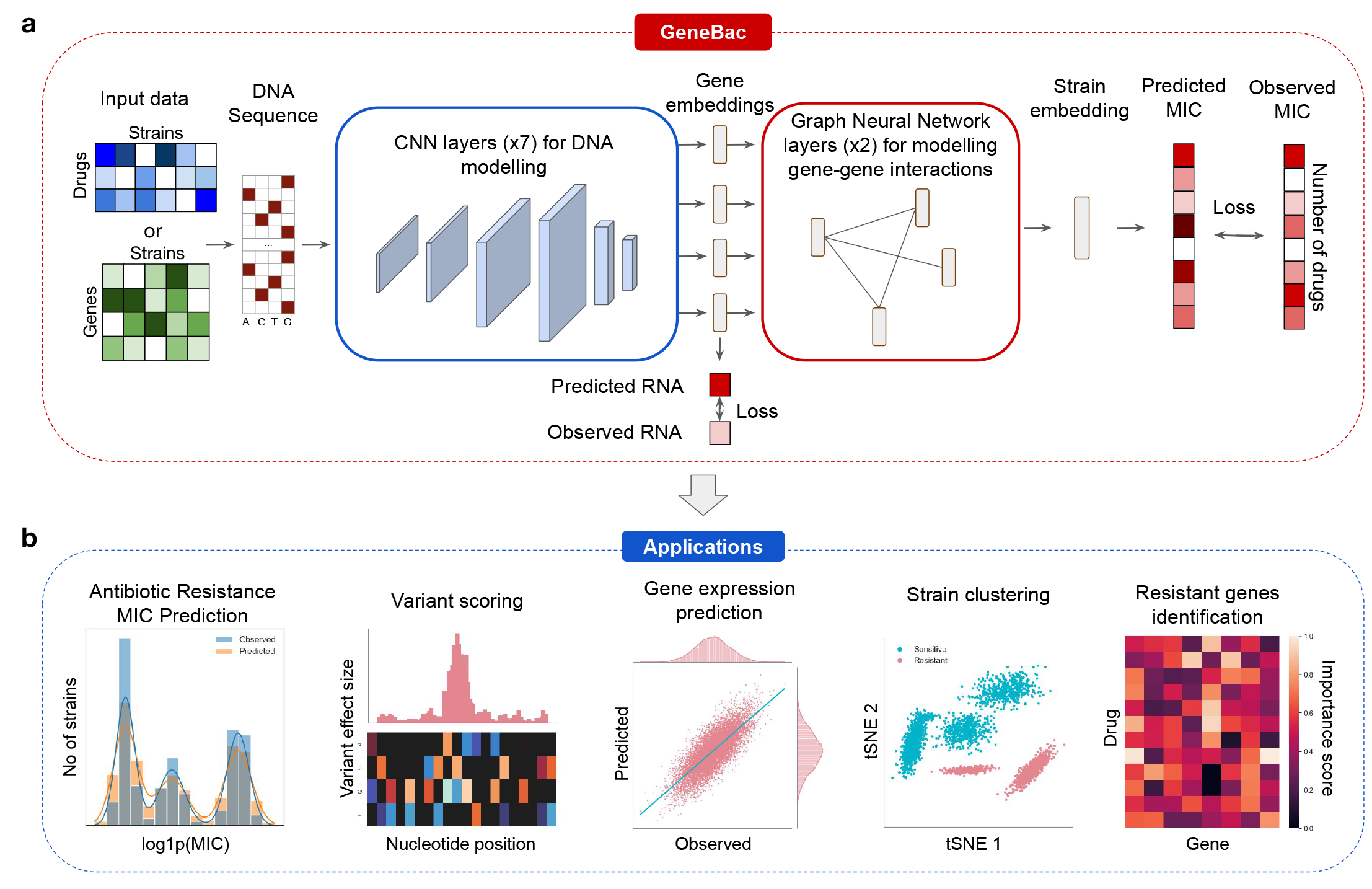
Overview of the GeneBac framework and its applications. *(a)* Given DNA sequence of a bacterial strain and appropriate labels, GeneBac can be trained to predict MIC for various drugs or RNA abundance. It takes as input one-hot encoded DNA sequence from a single strain with the variants integrated into a sequence. The input is processed by a CNN to extract features from DNA and a GNN to account for interactions between genes. *(b)* We apply GeneBac to a number of tasks including antibiotic resistance prediction, variant effect prediction, gene expression prediction, strain clustering and resistant genes identification.

We apply GeneBac to *Mycobacterium tuberculosis* and *Pseudomonas aeruginosa* datasets. GeneBac accurately predicts the MIC across two different bacterial species and substantially outperforms existing methods on antibiotic resistance prediction across different drugs. We demonstrate that GeneBac is a modular framework that can be applied to a diverse set of tasks including antibiotic resistance, variant effect and gene expression prediction as well as strain clustering and resistant genes identification (Fig. 1b). Finally, we show that GeneBac can utilize gene expression prediction task to learn transferable DNA features that can further boost the performance on the MIC prediction problem (Fig. 5).

## 2 Results

### 2.1 Overview of the GeneBac framework

We developed GeneBac, a framework for the prediction of antibiotic resistance from DNA sequences in bacteria. GeneBac allows for precise modelling of the genetic information of a bacterial strain by combining a Convolutional Neural Network (CNN), which captures the variation in the DNA, with a Graph Neural Network (GNN) designed to account for the interactions between different genes.

GeneBac takes as input multiple genes from a bacterial genome, which were pre-selected before training (Methods), with the variants present in the strain directly integrated into the input DNA sequences. Therefore, GeneBac allows to account for both single nucleotide polymorphisms (SNPs) as well as insertions or deletions (INDELs) of variable lengths in their genetic context. Each gene is represented as a one-hot encoded DNA sequence and consists of its protein-coding region as well as the region upstream of the gene. This way, we can account for variants in both the non-coding as well the coding part of the gene. Each gene is processed by a multi-layer CNN, without having access to the other genes at this stage. The CNN weights are shared across genes, which acts as a regularisation method and leads to learning common features across genes such as binding motifs and codons. The CNN processes each DNA sequence to a low-dimensional vector, representing the variation in the sequence of the gene.

The learned gene representations are then fed into the GNN as node features, where the edges between genes represent the interactions between them (Supp. Fig. 5a). Finally, the output of the GNN is used to predict the resistance to multiple drugs simultaneously. GeneBac is trained in the multi-task setup [24, 25], which allowed us to maintain a single model, instead of having a separate GeneBac model for every drug. In this way, representations are shared across various drugs. The GNN allows us to account for the dynamic interplay between genes and proteins and has led to improved performance across drugs (Supp. Fig. 2e) as well as increased model interpretability (Supp. Fig. 5).

GeneBac learns to accurately predict the MIC from the DNA sequence, and thus, can be used for scoring genetic variants. GeneBac also produces representations which could be used for other tasks such as gene expression prediction, strain clustering and resistant genes identification (Fig. 1b).

### 2.2 GeneBac predicts antibiotic resistance to multiple drugs

The MIC distribution of antibiotics often does not follow the binary pattern (Supp. Fig. 1a-b). However, existing methods treat the antibiotic resistance problem as a binary classification task [7, 12, 15, 22, 26, 27]. This can lead to inappropriate diagnosis in the clinical setting [17, 18] and makes it difficult to apply a trained model on a different dataset, as the binary threshold is usually dataset- and drug-specific. GeneBac solves this problem by directly predicting the MIC and solving a regression task, rather than predicting a binary value.

To benchmark our approach, we applied GeneBac to two datasets of clinical isolates: the *Mycobacterium tuberculosis* CRyPTIC dataset [28] containing the MIC readings for 13 different drugs, and a *Pseudomonas aeruginosa* (PA) dataset consisting of MIC data for 14 antibiotics [29]. The drug MIC values available vary strongly across strains (Supp. Fig. 1c), which has a strong impact on the variability of the results across drugs. The overall setup of the antibiotic resistance prediction task with GeneBac is shown on Fig. 2a.

**Fig. 2.**
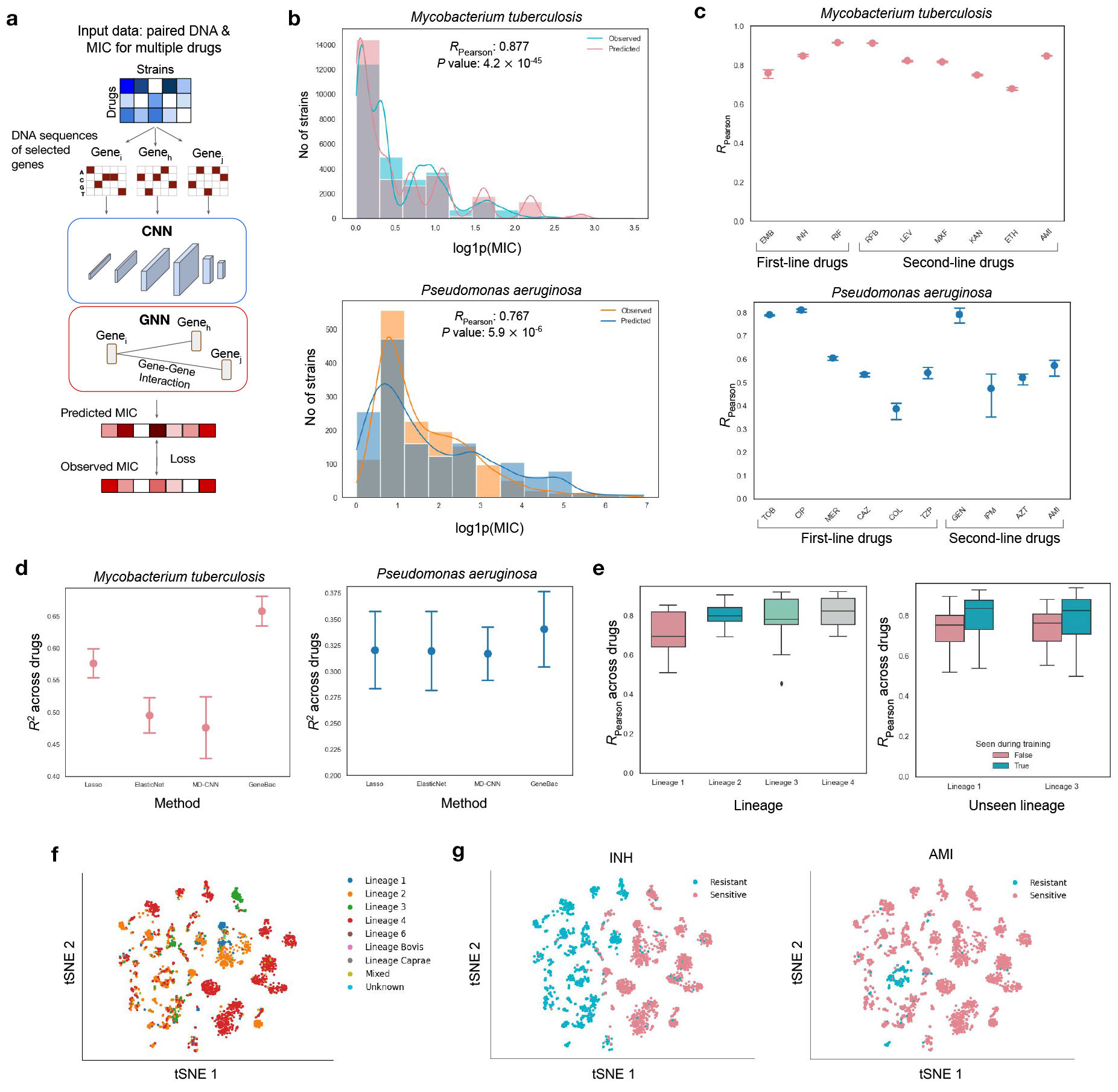
GeneBac antibiotic resistance prediction performance. *(a)* For antibiotic resistance prediction, GeneBac takes as input DNA sequences of multiple genes from a single strain. Each gene is processed independently by the CNN which converts it into a low-dimensional vector. The gene representation is used as node features in the GNN, which models the interactions between different genes. The final output of the GNN is used to predict the MIC to various drugs. *(b)* GeneBac observed and predicted MIC distribution across drugs and strains on held-out strains. *(c)* GeneBac prediction performance across first- and second-line drugs. The error bars represent the variance across runs and the dot represents mean performance. *(d)* Performance comparison of GeneBac and other methods on the antibiotic resistance MIC prediction regression task on first- and second-line drugs. The computed performance is the mean value across drugs and runs. The error bars represent variance across runs. *(e)* Performance across different lineages on the held-out strains from the MTB dataset (left). Results of the lineage ablation experiment on the held-out strains. The performance comparison is conducted between the lineage absent from training and the aggregate performance of other lineages seen during training. Each box represents the performance across drugs, with the horizontal line representing median performance. *(f)* Strain embeddings learned on the MTB dataset, coloured by lineage. *(g)* Strain embeddings learned on the MTB dataset coloured by their resistance to Isoniazid (left) and Amikacin (right).

First, we investigated how well can GeneBac predict the MIC on the held-out strains. GeneBac achieved high performance on both datasets, attaining over 0.87

Pearson correlation on MTB and 0.76 on PA across genes and strains (Fig. 2b). Furthermore, GeneBac performed well across first- and second-line drugs on both species (Fig. 2c), demonstrating the robustness of the method. We hypothesize that the difference in performance between MTB and PA datasets is due to the difference in the number of available training samples. Namely, the MTB dataset consists of 12,460 unique strains, while the PA dataset has only 1109 strains. To test this, we conducted a study where we subsampled the training data on the MTB dataset. Indeed, by doubling the number of training samples, the model performance increased by almost 30% including 47% improvement on the non-first-line drugs (Supp. Fig. 2b-c). This shows that further improvements in performance can be obtained by increasing the training set size.

GeneBac MIC predictions can be used for diagnosis by analyzing the MIC distribution of a drug. To demonstrate that, we divided GeneBac predictions into 3 separate categories: susceptible, intermediate and resistant. We then used the observed and predicted MIC to assign each observed and predicted value to a class and compute the accuracy per category (Fig. 2a). This has shown how the GeneBac MIC predictions can be used for picking appropriate antibiotic treatment while accounting for variable MIC distributions. Moreover, as there is no dataset-specific threshold, one could potentially use the MIC prediction of GeneBac on a strain from a different dataset.

A key challenge for computational methods for bacteria is accounting for the clonal population structure, which often results in the model outputting spurious findings [30]. This is a major issue for existing methods, which traditionally use as input a vector representing the presence of a variant in a strain [3, 7, 12–16]. In contrast, GeneBac uses DNA as input, learning features of the genetic code. Because of this, GeneBac is more robust to the variable structure of the bacterial population. We evaluated this by computing the performance per lineage on the held-out strains, where each lineage represents a population derived from the phylogenetic tree characterised by a similar set of variants. GeneBac performed well across all lineages (Fig. 2e). Moreover, to test whether GeneBac can generalise to new lineages, we completely removed strains of a certain lineage from the training set so that the model only saw it during testing. We performed it for two lineages and compared the results on the held-out strains from the unseen lineage with the performance across lineages which have been seen during training. We show that GeneBac can generalise to previously unseen linages, recording only a small decrease in performance between the unseen and seen lineages (Fig. 2e).

We further compared GeneBac against existing methods by reporting the average performance across drugs on the test set on the MTB and PA datasets (Fig. 2f). While we compared GeneBac to methods such as Lasso [31] and ElasticNet [32], we highlight that these methods are not capable of generalising to unseen variants, which is a significant limitation. However, we compared them to GeneBac in a simpler setting where this is not required. On the MIC prediction regression task, GeneBac achieved a mean *R*^2^ score across drugs of 0.66, against 0.48 achieved by MD-CNN and 0.58 by Lasso on the MTB dataset. On the PA dataset, GeneBac attained 0.34 *R*^2^ compared to 0.32 by MD-CNN and Lasso (Fig. 2d).

We also computed performance on the binary task (Supp. Fig. 3), with GeneBac outperforming other methods on the area under the receiver operating characteristic (AUROC) on both datasets (Supp. Fig. 3a). Furthermore, GeneBac achieved the most balanced results on specificity and sensitivity across drugs, with other models recording high specificity, but low sensitivity (Supp. Fig. 3b).

In addition to predicting antibiotic resistance, GeneBac also produces strain representations, which can be used for clustering and investigating strain populations. We extracted low-dimensional strain embeddings using the trained model and visualised them in two dimensions using the t-distributed stochastic neighbour embedding (t-SNE) [33]. We observed that across different datasets and drugs, the resistant and susceptible strains tend to cluster together (Fig. 2g & Supp. Fig. 6). Moreover, we also visualised the MTB strains by their lineage (Fig. 2f), noting that strains of the same lineage tend to cluster together. This shows that GeneBac can learn the population structure while not explicitly accounting for it. Additionally, while many strains clustered by their lineage, we observed that highly resistant strains concentrate in the same neighbourhood, despite their lineage. This is expected as drug resistance does not need to be lineage-specific.

### 2.3 GeneBac predicts the effect of seen and unseen variants

A key limitation of existing methods is their inability to account for variants that were not observed during training. This is particularly important in the diagnosis setting, where new variants are almost guaranteed to appear considering the rapid rate of mutation in bacteria. As a result of using DNA with variants integrated into the sequence as input, GeneBac is uniquely capable of accurately predicting the effects of specific variants, including new ones. To test this claim, we extracted and scored the variants present in the WHO MTB catalogue of mutations associated with drug resistance [4] with GeneBac trained on the MTB dataset.

We performed in silico mutagenesis (ISM) for each variant and recorded the variant effect size measured as the delta between the input with and without the variant. GeneBac was able to recover a number of variants that have been shown to lead to increased resistance, such as the non-coding transcript variant A1401G located in the rrs gene, missense variant H445D in the rpoB gene (Fig. 3d) and deletion katG 1351 in the katG gene. The mean variant effect size is 0.75 for variants associated with resistance, compared to 3.7 10*^−^*^3^ for variants not associated with resistance and 10*^−^*^4^ for synonymous variants (Supp. Fig. 4b). Importantly, the model performs well on predicting variant effect size on variants not seen during training (Fig. 3b), showing that the model can generalise to new variants. GeneBac can also predict the effect of variants across distinct drugs. We computed the performance for a set of variants from the catalogue associated with different drugs. GeneBac successfully predicted the variant effect size across multiple drugs (Fig. 3c), showing that it can be applied to a range of first- and second-line drugs. We performed a similar analysis across genes, demonstrating that GeneBac accurately predicts variant effect size on distinct loci and avoids relying on a specific gene (Supp. Fig. 4c).

**Fig. 3.**
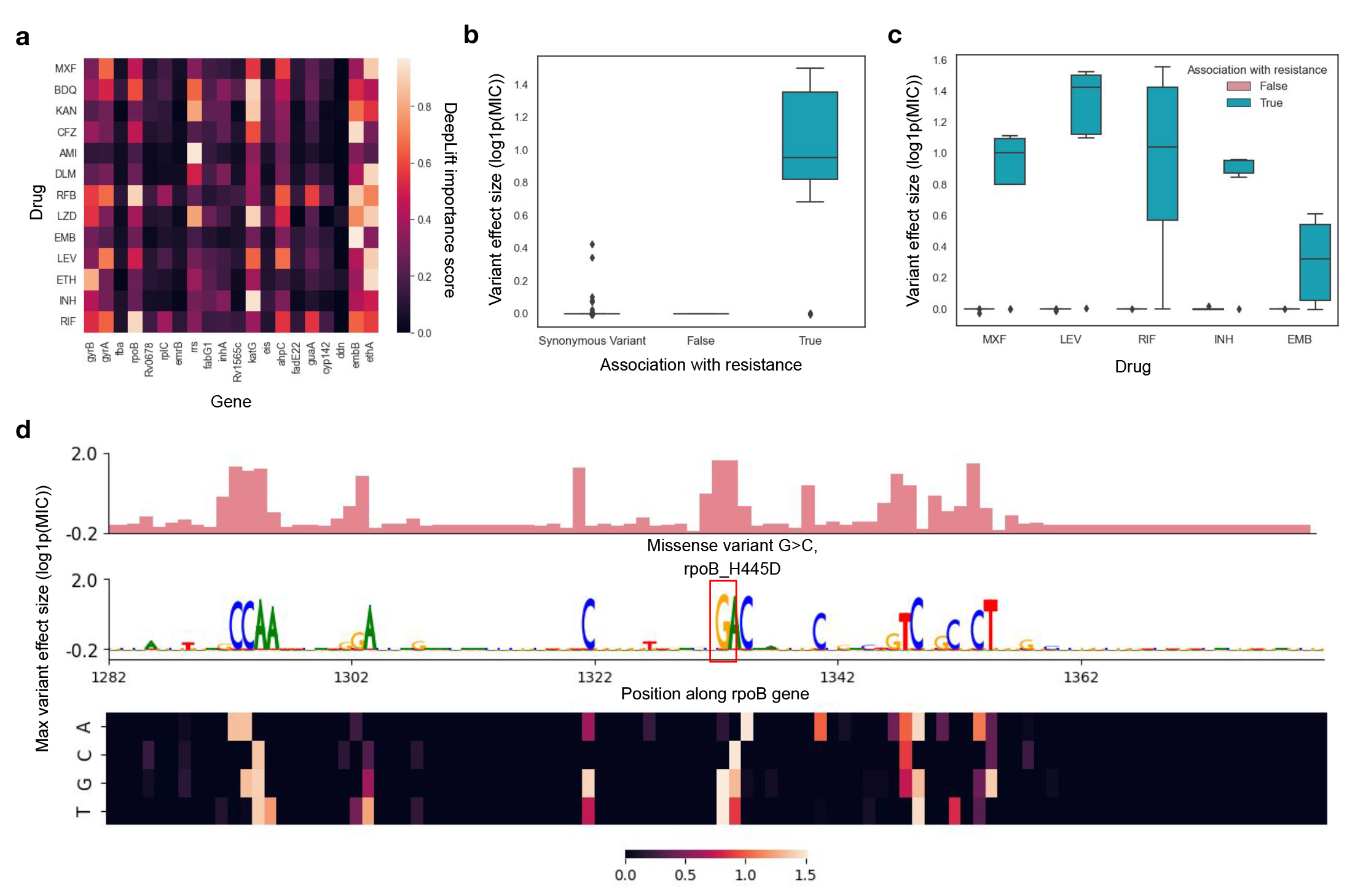
GeneBac predicts variant effect. *(a)* Drug-Target importance scores aggregated across held-out strains and normalised per target. *(b)* Variant effect size of novel variants unseen during training from WHO MTB mutation catalogue across variants. The variant effect size was calculated using the GeneBac model trained on the MIC regression prediction task on the MTB dataset. Each box represents the variant effect size across different variants with the horizontal line representing the median variant effect size. *(c)* Variant effect size of variants from WHO MTB mutation catalogue across different drugs on variants associated and not associated with resistance. Each box represents the variant effect size across different variants with the horizontal line representing the median variant effect size. *(d)* GeneBac predicts the effect of the variant rpoB H445D associated with the resistance to Rifampicin (RIF). The variant was not seen during training. The plot shows the results of in silico mutagenesis of the region from 1282 to 1382 downstream of the start codon of the rpoB gene. The top and middle plots show the nucleotide with the highest variant effect size, while the bottom plot shows the variant effect size per nucleotide position.

By leveraging DNA as input, GeneBac can predict the effects of variants in non-coding as well as coding sequence. We demonstrated this by dividing the variants associated with resistance from the catalogue into 3 distinct types, (1) missense variants, (2) insertions or deletions (INDELs) and (3) non-coding variants, and computing the variant effect size across these types. GeneBac performed similarly well across the variant types (Supp. Fig. 4d), showing that it can model different kinds of mutations Identifying drug-resistance genes can help understand the molecular mechanisms leading to resistance, thus supporting the development and repurposing of existing antimicrobials. Motivated by this, we used GeneBac to evaluate whether it can recover the genes associated with resistance to particular drugs. To achieve this, we computed and aggregated at the gene level the importance scores per drug using DeepLift [34].

Fig. 3a shows the normalised importance score for each combination of drug and gene. GeneBac recovered a number of genes which have been shown to be associated with resistance to drugs. This includes rrs for Amikacin [35], embB for Ethambutol [36], katG for Isoniazid [37]. For PA, GeneBac also highlighted a number of drug-resistance genes. Such as gyrA, which is an important gene across multiple drugs, which we have corroborated with existing literature [38, 39] and parC for Ciprofloxacin [40] (Supp. Fig. 4a). Moreover, GeneBac also highlighted a number of drug-resistance genes which have not been previously reported.

Finally, leveraging the Graph Neural Network allows us for a more interpretable introspection of the model prediction. Using GNN Explainer [41] to identify the most important nodes and edges on a set of hold-out strains, we showed how the GNN can be used for analysis at the strain level, helping to identify key genes and their interactions for the final prediction (Supp. Fig. 5c-f).

### 2.4 GeneBac predicts gene expression

Accurate prediction of the phenotype, such as antibiotic resistance, often requires understanding how the variation in the DNA sequence affects gene transcription. To validate that GeneBac can capture the effect of genetic variants on transcription, we used the CNN employed in GeneBac for RNA abundance prediction. We trained the CNN (Fig. 4a) on the dataset of 386 PA strains containing paired DNA and RNA readings [42]. From each strain, we extracted the genes present in the core genome amounting to 4,548 genes. To learn common features across the genome, we treated each gene in a strain as a single example, resulting in over 1.75 million (400 4, 548) samples.

**Fig. 4.**
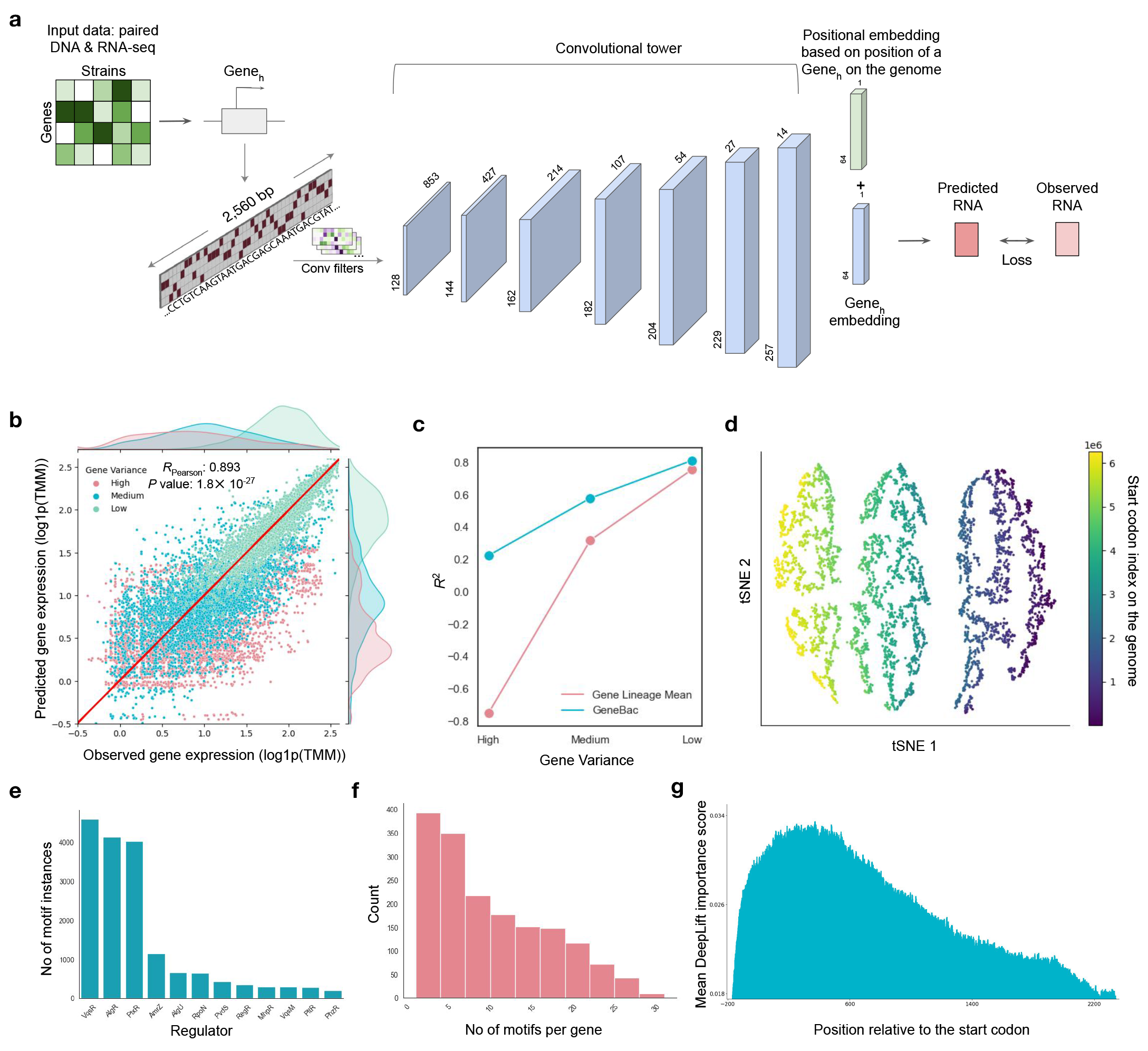
GeneBac predicts gene expression. *(a)* GeneBac can be trained to predict the RNA abundance from the DNA sequence of a gene. The one-hot encoded DNA sequence of a gene is processed by the GeneBac CNN module which outputs a gene embedding. The gene embedding is combined with the positional gene embedding based on the position of the start codon in the genome and used to predict the RNA abundance. The architecture of the CNN module is the same across antibiotic resistance and gene expression tasks. The architecture visualisation was adapted from the scBasset model [22]. *(b)* GeneBac observed versus predicted gene expression on the PA dataset coloured by category of gene variance. For visualisation purposes, we randomly sampled examples to maintain a similar number of samples across the gene variance category. *(c)* Performance comparison of GeneBac versus the mean expression of the gene in the lineage across gene variance categories on the held-out strains. The performance is computed as the mean value across genes and strains. *(d)* t-SNE visualisation of the gene embeddings learned by the CNN final layer on the PA RNA-seq dataset, coloured by the location of the start codon on the genome. *(e)* Number of motif instances per regulator found on the PA reference genome. *(f)* Number of binding motifs in genes with one or more detected binding motifs found on the PA reference genome. *(g)* The mean importance of each base-pair position across genes on the PA reference genome. The results were normalised to account for various gene lengths.

**Fig. 5.**
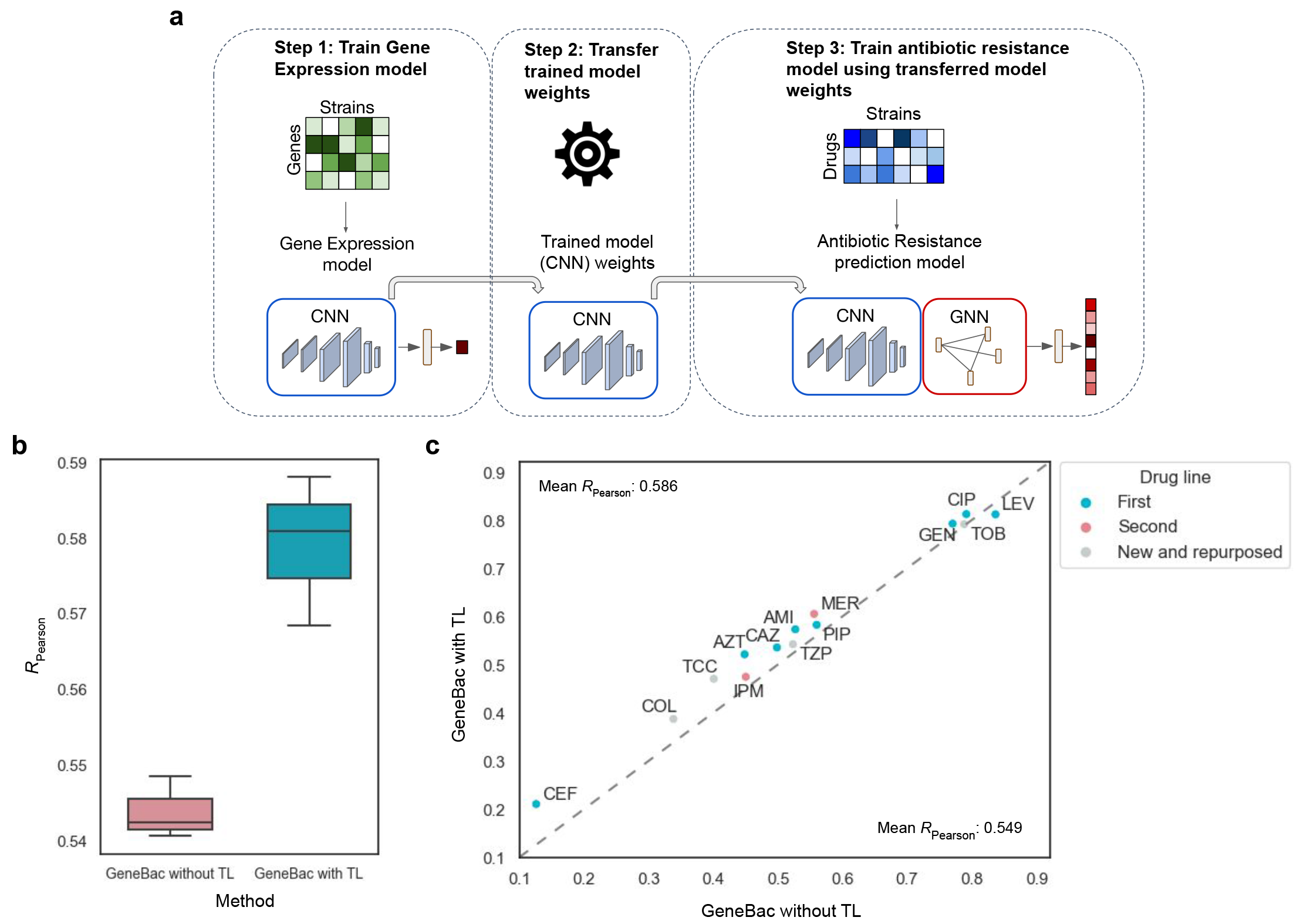
GeneBac enables transfer learning across modalities. *(a)* The schematic of the transfer learning workflow. *(b)* Performance comparison of the GeneBac model with and without transfer learning (TL) across numerous runs. Each box represents the mean drug performance variance across runs, with the horizontal line representing median performance. *(c)*, Pairwise per drug performance comparison of GeneBac model with and without TL, coloured by drug line.

We trained the model to predict the *log*1*p* normalised RNA abundance and tested it on the set of held-out strains. GeneBac accurately predicted gene expression, achieving almost 0.9 Pearson correlation across drugs and strains (Fig. 4b). Based on a standard deviation of expression across strains, we divided the genes into 3 categories with low, medium and high variance and computed performance across them (Fig. 4c). We benchmarked our results against the mean expression of the gene in the lineage, showing that GeneBac significantly outperforms it on genes with high and medium variance. The gap between the two is much smaller for low-variance genes, which is expected as a large number of these genes are housekeeping genes with very little variation in expression across strains.

CNN models have been shown to be able to extract patterns from the DNA sequence such as transcription factor binding motifs [21, 22]. To validate that our model is able to do that, we used TF-MoDISco [43] to discover motifs extracted by our model based on the DeepLift [34] importance scores coming from the trained model. GeneBac identified a number of regulators (Fig. 4e) in various regions (Fig. 4f). Discov- ered regulators include a key quorum-sensing regulator VqsR [44, 45], a transcriptional regulator involved in pathogenesis AlgR [46] and sigma factor AlgU [47].

To investigate the nucleotide positions that are most influential for predicting RNA abundance, we ran the DeepLift [34] algorithm on the reference genome and computed the average importance score per nucleotide position across genes, normalised by gene length. The important scores show significant enrichment around the start codon and downstream of it, within the body region, which aligns with the reported TF binding sites in PA [48, 49].

Bacterial genes are known to cluster by their position on the genome, often sharing similar functions and forming operons [50, 51]. To examine whether gene representations outputted by GeneBac cluster by their position on the genome, we extracted the gene embeddings from the trained gene expression model. We then visualised the gene embeddings using the t-SNE algorithm, showing that genes cluster by their position on the genome (Fig. 4d).

### 2.5 GeneBac transfer learns across modalities

Many bacterial genomics datasets consist of only a couple of hundred samples, making it difficult to efficiently apply more complex machine learning models, which require a large number of samples. GeneBac is a modular framework, which means that it can be applied to several tasks, including gene expression and antibiotic resistance MIC prediction. We leveraged this modularity and used transfer learning (TL) to improve the performance on the MIC prediction task by using GeneBac which was initially trained on the gene expression dataset (Fig. 5). Specifically, we firstly trained the CNN model on the PA gene expression dataset described in the previous section and then used the trained weights to initialise the CNN part of the GeneBac for the task of antibiotic resistance prediction in PA (Fig. 5a). The motivation behind this is that as the input to the model is in both cases DNA sequence, the model can leverage the features learned in the gene expression problem, such as binding motifs, to improve performance on the MIC prediction task.

Transfer learning across modalities improved mean performance across drugs by almost 4% as measured by Pearson correlation (Fig. 5b). Moreover, TL boosted performance on 13 out of 14 drugs on Pearson correlation, particularly improving the results on drugs with lower general performance (Fig. 5c), which often have fewer training samples available (Supp. Fig. 1c). TL increased the Pearson correlation coefficient by 8% on the second-line drugs and 9% on new and repurposed drugs, compared to 5% improvement on first-line drugs.

## 3 Discussion

The emergence of bacteria resistant to antibiotics and other treatments is a global threat, making tools that can rapidly diagnose patients as well as help us understand the molecular mechanisms of AMR crucial for global health. In this study, we present GeneBac, a deep learning framework for predicting the minimum inhibitory concentration (MIC) of antibiotics from DNA sequence by integrating interactions between genes.

Two properties make GeneBac a unique tool in the bacterial genomics toolbox: (1) the ability to directly predict the minimum inhibitory concentration (MIC) of multiple drugs, and (2) leveraging genetic context which enables GeneBac to generalise to previously not observed variants and make it more robust to the population structure. These properties translated to improved models for two key problems of biological relevance: antibiotic resistance MIC and variant effect prediction. We demonstrated that GeneBac can accurately predict the MIC across various drugs and distinct bacterial species, as well as RNA abundance. These results suggest that the model can learn the impact of the variation in the DNA on antibiotic resistance and gene expression. Using GeneBac, we can predict the effect of an arbitrary variant, including novel variants which have not been seen in the training data, and employ it for identifying variants which cause a notable change in the MIC of an antibiotic. Furthermore, GeneBac can also score genes, facilitating antibiotic-resistant gene identification. GeneBac is a modular framework relying on the DNA sequence as input. We used it to improve the performance on the MIC prediction task, by transfer learning from the gene expression task, with both tasks using DNA as input.

Several paths for further improving our framework appear promising. The performance of machine learning models, including GeneBac, depends on the training data. Therefore, curating data, as well as leveraging cross-species and cross-modality data through transfer learning would likely boost performance. Specifically, it would be beneficial to curate such large-scale datasets for bacterial species whose genome is not as stable as in *Mycobacterium tuberculosis*. Recent work demonstrated the capability to predict the effect of a mutation in a protein sequence on its fitness [52]. Combining such models with GeneBac which works on the DNA sequence level could improve its performance on the MIC prediction and provide more interpretable results.

With the development of larger datasets, we envision that GeneBac could be systematically employed for diagnosis in the clinical setting and uncovering causal variants as well as genes influencing antibiotic resistance. To foster these applications, we made the model openly available along with checkpoints and tutorials. Furthermore, we precomputed a set of variants in *Mycobacterium tuberculosis* which have shown a significant effect on the MIC for each drug. We hope that our model will facilitate the development of improved diagnostic tools and stimulate the understanding of the molecular mechanisms leading to the antibiotic resistance.

## 4 Methods

### 4.1 Data processing

#### Antibotic Resistance MIC processing

We downloaded the CRyPTIC *Mycobacterium tuberculosis* [28] dataset from http://ftp.ebi.ac.uk/pub/databases/cryptic/release june2022/reproducibility/

data tables/cryptic-analysis-group/. This included an already processed variant matrix as well as MIC values per strain. We excluded the strains with “Low” quality MIC readings. For the regression task, we normalised the data with *log*1*p*, while for the binary task, we used labels already provided in the dataset. We used the H37Rv reference genome and accompanying genome assembly downloaded from https://www.ncbi.nlm.nih.gov/nuccore/NC 000962.3 to create input data by integrating it with the variants present in the variants matrix. As the available MIC readings vary per strain, we replace the values where the MIC of a drug is missing with 100, which we later use in the model to exclude from loss and metrics computation. We visualise the per-drug MIC distribution as well as the number of samples with MIC available per drug in the Supp. Fig. 1c). We make the processed data as well as the preprocessing script available as part of the code for reproducibility.

We use *Pseudomonas aeruginosa* dataset. We follow the preprocessing steps at the CRyPTIC data mentioned above, using the reference genome PAO1 together with its genome assembly from https://www.ncbi.nlm.nih.gov/nuccore/NC 002516.2. As this data does not come with preassigned labels, we manually analyse and set the binary threshold for the binary task of antibiotic resistance prediction based on the MIC distribution of a drug.

#### RNA-seq processing

For the RNA-seq data used for the gene expression task, we used the PA dataset of 386 strains from https://www.ncbi.nlm.nih.gov/geo/query/acc.cgi?acc=GSE123544 [42]. The data consists of paired DNA-sequence and RNA counts readings. We use only the core genome amounting to 4,548 genes. We follow standard preprocessing steps by normalising the gene expression data by performing TMM and *log*1*p* normali- sation [53]. For the accompanying DNA sequence data, we leverage the reference genome PAO1 together with its genome assembly from https://www.ncbi.nlm.nih.gov/ nuccore/NC 002516.2, integrating the called variants directly into the DNA sequence, same as with the DNA sequences used for MIC prediction.

#### Selecting input genes

For the antibiotic resistance task, taking in a full bacterial genome, which in the case of the MTB and PA amounts to approximately 4.5 and 6.3 megabase respectively is both computationally challenging and can easily lead to overfitting due to the large length of the genome compared to the number of samples. Due to this, we preselect a set of genes which we use as input to the antibiotic resistance models.

We collated the genes from the literature, including the analysis of results of existing methods such as GWAS to choose a set of suitable genes covering the available drug. We also tried training the model with different sets of preselected genes, including most variable genes, defined as the number of variants per gene across strains normalised by gene length, choosing the optimal set of genes based on cross-validation results. We include the table with the complete set of genes used for MTB in Supp. Table 1. The table includes the gene, its location and its associations to drugs available in the dataset.

Having to preselect genes is a major limitation, however, it is a major step forward compared to methods that require preselecting the variants for antibiotic resistance prediction. Therefore, we consider having to preselect genes, rather than specific variants as an improvement. We believe that with improved architectures, in the future we will be able to expand to cover a much larger part of the genome.

#### DNA sequence processing

A large majority of the existing DNA sequence methods [19, 21–23] leverage solely the reference genome as the input DNA sequence as the data used in such methods lack information on the variant present in the DNA. In this work, we directly integrate the variants present in the strain, therefore each strain has a unique DNA sequence. To get input data, we start from the reference genome together with the accompanying genome assembly. We then use the variant matrix to directly annotate the variants present in a strain, accounting for both single nucleotide polymorphisms as well as INDELs. As part of the script, we run sanity checks to ensure the resulting genomes are correct.

The output of the processing script is a DNA sequence for each gene for all strains. As part of the gene, we include the region upstream of the start codon, which is 200 bps for PA and 100 bps for MTB. This is so as to account for variants in both coding and non-coding sequence. We set the maximum length of the gene at 2,560 bps which fully covers more than 96% of genes in both MTB and PA. We pad the gene sequence if it is less than 2,560 bps.

### 4.2 Modelling

#### Model architecture

GeneBac is a neural network architecture that can be used for the prediction of (1) antibiotic resistance of a set of drugs for a bacterial strain, or (2) RNA abundance. For the first application, which is our main use case, GeneBac takes as input a set of DNA sequences, each representing a different gene with the present variants directly integrated into a sequence and prior information on the interactions between considered genes. In the case of the RNA abundance prediction, the model takes as input a single gene. Each gene consists of its region upstream and the protein-coding region. The DNA sequence of a gene is then one-hot encoded resulting in a 2, 560 x 4 matrix used as input to the CNN.

The model consists of 2 modules, (1) a CNN taking as input the DNA sequence and outputting a low-dimensional representation of a gene, and (2) a GAT leveraging the gene representations as node features and modelling the interactions between the genes using data from existing protein-protein interaction database [54]. As bacteria show spatial gene expression patterns, with the location of the gene on the genome from the origin of the replication affecting its mean expression [55, 56] and genes often clustering together in operons [50, 51], we inject positional information by adding absolute (i.e. not relative) positional encodings to the gene embedding.

##### Convolutional Neural Network

The CNN takes as input DNA sequence and outputs low-dimensional gene representations. It consists of:

- 1D convolutional layer with 128 filters of size 17 x 4, followed by batch normalization, GELU and width of 3 max-pooling layers, resulting in the output matrix of dimension 128 x 853.
- A series of 6 convolutional blocks, also referred to as the convolutional tower, consisting of batch normalisation, max pooling and GELU layers. We increase the number of filters in each layer (144, 162, 182, 204, 229, 257). Each layer has a kernel width of 6 and is followed by a max pooling layer with a width equal to 2. The output of the convolutional tower is a matrix of dimension 257 x 14. Such architecture leveraging a series of layers with an increasing number of filters has been shown to perform well on the DNA modelling task [19, 22].
- A fully connected layer converts the output to a vector of 64 dimensions, followed by batch normalisation, a dropout rate of 0.2 and a ReLU activation function.

In the case of antibiotic resistance prediction, the CNN module takes as input multiple distinct genes and processes them independently, with the same weights being used for each gene. The gene representations are then used as node features for the graph model. For the gene expression prediction task, the input is the DNA sequence of a single gene. The low-dimensional gene representation is then fed into a fully connected layer with an output of 1, representing the predicted *log*1*p* normalised RNA abundance.

##### Graph Neural Network

The Graph Attention Network (GAT) [57], uses gene representations from the CNN as node features and prior information from the STRING database [24] to construct the graph and extract the edge features. Using the selected genes in MTB or PA, we query the STRING database setting a confidence threshold at 0.4, which is the default and use the resulting edges between genes to construct the graph. We also include a vector of edge features for each edge. Such a strain-level graph is then fed into:

- A two-layer graph attention network, with the input, hidden and output dimensionality in each graph layer equal to the size of gene representation outputted by the CNN. Each graph layer is followed by a dropout of 0.2 and a Leaky ReLU activation function.
- The output of the graph layers is then flattened and passed through a fully connected neural network converting it into a strain embedding, designed to serve as a low dimensional strain representation. This is followed by a dropout of 0.2 and the ReLU activation function.
- Finally, this strain representation is fed into a fully connected layer with an output equal to the number of drugs available in the dataset.

##### Positional embedding

Bacterial gene expression depends on the position in the genome [55, 56]. To account for this, we add fixed absolute positional embedding to the gene representation, right after the CNN and before the graph module (Fig. 4a). The dimensionality of positional embedding is equal to the gene representation dimensionality. Given *t_i_*, which denotes the position of the start codon of the gene *i* on the reference genome, a coefficient *j* and a scaling factor *s*, the positional embedding is given as:

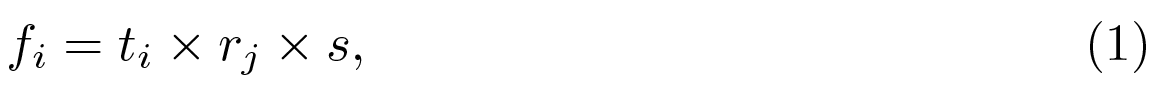

where *r_j_* is placed linearly between the coefficient *j* and its *log*. We set the coefficient *j* to 3.65, based on bacteria’s spatial gene expression pattern [56] and the scaling factor *s* to 10*^−^*^7^.

#### Training approach

We trained the antibiotic resistance MIC prediction and the gene expression models, both regression tasks with the mean squared error (MSE) loss and optimised the model for the *R*^2^ score, also known as the coefficient of determination. The binary antibiotic resistance model was trained with binary cross-entropy loss and optimised for the area under the ROC curve (AUROC).

For all antibiotic resistance models, we split the data by strain and performed 5- fold cross-validation to choose optimal hyperparameters. We trained the model for a maximum of 500 epochs on both the MTB and PA datasets. After selecting the optimal hyperparameters, we optimised the antibiotic resistance regression models for the mean train *R*^2^ score across drugs and the mean train AUROC score across drugs for the binary ones. For the gene expression model, we also split the data by strain, but because of the larger data size amounting to almost 2 million samples, we opted for training, validation and test split and optimised the model for validation *R*^2^, training the model for a maximum of 100 epochs. We experimented with different numbers of filters in the CNN, the number of graph layers in the GAT, maximum gene length, different sizes of the gene and strain embeddings including as well as excluding positional embeddings. The full table of optimal hyperparameters for all models can be found in Supp. Table 2 & 3.

As in the antibiotic resistance prediction task, the MIC readings available vary strongly per strain, we ignore the drugs with MIC not available, only backpropagating through the drugs for which we have available data in the strain.

All training runs were performed on a single GPU and the model parameters were updated with stochastic gradient descent using the Adam optimiser with the gradient clipping rate set to 1. The hyperparameters were tuned using grid search. The whole antibiotic resistance model has 1.5 million parameters, which is significantly less than MD-CNN [27], which has more than 60M parameters using the same input.

## 5 Code Availability

Code for training and using the model including checkpoints and instructions how to run the model for different applications can be found at: https://github.com/ macwiatrak/GeneBac.

## 6 Author Contributions

M. W. conceived the idea and contributed to the theoretical and experimental design.

M. W. contributed to data preprocessing. M.W. coded up the model and performed the experiments.

A. W. contributed to the data acquisition, data preprocessing, experimental design as well as writing of the manuscript. A. D. contributed to the data acquisition, data preprocessing and experimental design.

A. F. and M. B. contributed to the experimental design and writing of the manuscript.

## Acknowledgements

We would like to thank Vitalii Kleshchevnikov, Dino Oglic, Alexander Predeus, Jacob Hepkema and Mohammad Lotfollahi for helpful discussions and Lorenz Kretschmer for feedback on the manuscript.

## Supplementary Information

Below, we describe the details of the settings of the experiments conducted in the study. For implementation and further details, please see our github repository https://github.com/macwiatrak/GeneBac.

### Antibiotic resistance prediction

#### Generalising to unseen lineages

To test whether GeneBac is robust to the clonal population structure and thus, can generalise to unseen lineages, we completely removed a lineage from the training set, trained GeneBac on the new training set and evaluated the performance on the removed lineage.

We removed two lineages, one at a time, Lineage 1, which has 699 samples in total and Lineage 3 with 1069 samples. We redid the train test split so as the maintain the same proportion between train and test, however, we made sure that the removed lineage is only present in the test set. We then trained GeneBac model on such splits and compared the performance across drugs on the strains from the removed lineage versus the performance on the strains from other lineages, which were seen during training.

#### Benchmarking antibiotic resistance prediction methods

We benchmarked GeneBac against a set of existing methods. Firstly, we compared it against two well-known linear models, Lasso [31] and ElasticNet [32]. We chose them because they have been shown to perform well [7] and because they can limit the set of considered variants during training, thus avoiding the situation where the number of predictors (here, variants) exceeds the number of available samples. However, one key limitation of such models is that they are not able to generalise to unseen variants. Secondly, we benchmarked our model against MD-CNN [27], which is a also CNN- based model, but with a strongly different architecture. We did not consider methods that require pre-selecting variants as we believe that it’s a major limitation.

For both MIC and binary label prediction and all methods, we used the same input, output and data split. We run the models across multiple seeds and reported the performance across seeds.

##### Lasso & ElasticNet

We processed the variant matrix to extract a one-hot vector per strain, where each value in a vector represents the presence or absence of the mutation. We then used this vector as input to the model.

We used the Lasso & ElasticNet implementations from the *scikit-learn* package [58]. We trained the model separately for each drug. We also experimented with the multi-task learning setup, but it yielded worse results than the single-drug setup. For each drug, we tuned the hyperparameters, such as the *l*_1_ regularisation strength for Lasso and *l*_1_ ratio as well as *α* for ElasticNet using grid search and 5-fold cross-validation.

After tuning we trained the model on the training set and tested it on the held-out strains.

We followed the same approach for both the MIC prediction regression and binary label prediction tasks.

##### MD-CNN

We used the open-sourced implementation of MD-CNN [27] from https://github. com/aggreen/MTB-CNN to reimplement the model in PyTorch. We used the same DNA sequence input as for the GeneBac model, however, in contrast to GeneBac, we extended the maximum length of the gene to the longest gene in the input. We also ran MD-CNN with the same maximum length of the gene as for GeneBac, but it yielded worse results during the cross-validation.

We used the same data splits, including cross-validation as for the GeneBac and tuned the hyperparameters during 5-fold cross-validation. We trained MD-CNN in the multi-task setup and report the results on the held-out strains. We notice during cross-validation that MD-CNN quickly overfits. This is motivated by the large number of parameters of the MD-CNN model amounting to *>*60M parameters compared to GeneBac’s 1.5M parameters using the same input. By performing in silicon mutagenesis, we also noticed that MD-CNN tends to rely on a small subset of variants for the final prediction, thus hurting its generalisation capabilities.

We followed the same approach for both the MIC prediction regression and binary label prediction tasks.

#### Metrics

For the MIC prediction regression case, we used two metrics for evaluating our results, (1) the Pearson correlation coefficient, which measures the linear correlation between two sets of data and (2) the *R*^2^ score, also known as the coefficient of determination, which provides a measure of how well observed outcomes are replicated by the predictive model, based on the proportion of total variation of outcomes explained by the model. The *R*^2^ score is given by:

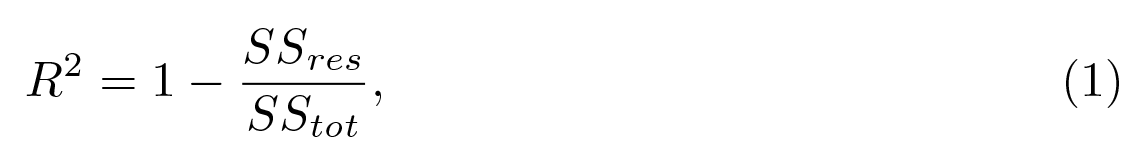

where 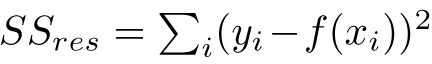 is the sum of residual squares, and 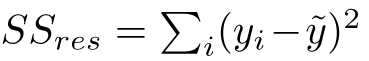 is total sum of squares.

For the binary antibiotic resistance prediction, we used the standard set of metrics used for evaluating antibiotic resistance models. This includes the area under the ROC curve (AUROC), which measures the model’s ability to discriminate between positive (resistant strains) and negative (susceptible strains) samples, specificity and sensitivity. Specificity and sensitivity are metrics used in the clinical setting to evaluate the model’s performance. Specificity gives the probability of a negative result (susceptible), conditioned on the sample being truly negative and is given as 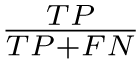 . Sensitivity is the probability of a positive result (resistant), conditioned on the sample being truly positive, hence it can be described as 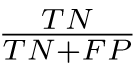 . Where, TP stands for True Positive, FN for False Negative, TN for True Negative and FP for False positive. To evaluate the balance between specificity and sensitivity, we computed their geometric mean. This is given as 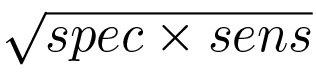.

We report the performance across drugs, as a measure of overall model performance, as well as per drug performance.

#### Strain clustering visualisation

To evaluate, whether trained GeneBac produces strain representations which cluster in a meaningful way. We extracted the low-dimensional (64) strain embeddings, which are produced by the layer preceding the final layer from the datasets. We then used t-distributed stochastic neighbour embedding (t-SNE) [33] to visualise the representations in two dimensions. In the t-SNE, we set perplexity to 100 and the number of iterations to 800 for MTB. For PA, we set perplexity to 40 and the number of iterations to 1000 for PA.

We visualised the resistance to antibiotics using the provided labels. We have visualised how strains cluster by lineage on the MTB dataset Fig. 2f. We did not do it for PA, as in the PA dataset there is no clear set of distinct lineages.

#### Variant effect prediction

##### Using WHO *Mycobacterium Tuberculosis* Mutation Catalogue

Using the WHO MTB mutation catalogue [4], we selected the variants present in the genomic loci used to train the model and associated with the drug present in the dataset, resulting in 3,009 unique variants. To ensure quality, we removed the variants present with the *interim* confidence grade. We divided the variants into those seen and unseen during training by overlapping them with the variants present in the variant matrix of the dataset. This provided us with 2412 unique variants, out of which 1057 were found to be present in the training set and 1355 were not present.

To assess the effect of the variant on the predicted MIC of a drug, we introduced the variant into the input DNA sequence and using GeneBac trained to predict MIC on MTB computed the predicted MIC values. To get the variant effect size, we took the delta between the MIC value for a drug of interest predicted using the DNA sequence with and without the variant. We compute such score per variant and use it for analysis.

##### Predicting variant effect across drugs & genes

To measure whether GeneBac can predict the variant effect across distinct drugs & genes. We divided the aggregated variants across drugs or genes and associated as well as not associated with resistance. To make the analysis meaningful, we exclude drugs which have less than 5 unique variants in each category (associated and not associated with resistance). We follow the same approach for the analysis across genes. We then use this data to visualise the results (Fig. 3c & Supp. Fig. 4c).

##### Predicting variant effect across mutation types

GeneBac takes as input the DNA sequence, therefore allowing to easily include arbitrary mutations, such as single-nucleotide polymorphisms (SNPs) as well as insertions or deletions (INDELs) in both the coding and non-coding region of the gene. To evaluate whether GeneBac can generalise to various mutations, we used the variant type in the catalogue denoted as the *effect* to group the mutations into 3 types, (1) missense variant, (2) non-coding variant and (3) INDEL. As there is only a small number of non-coding variants and INDELs which are associated with resistance in the catalogue (3 and 2 respectively), we included the variants with *interim* confidence grade. This included a lot of variants with low confidence, which impacted the results, however, it allowed us to observe the trend. We then computed the results across different variant types and their association with resistance. The results are visualised on the Supp. Fig. 4d.

##### Resistant genes identification

To identify genes which may be causing the resistance to given antibiotics, we used DeepLift [34] to run and collect input DNA importance. We then aggregated the absolute importance per base pair on the gene level for each drug. Finally, we used min-max normalisation to normalise the importance of each gene for a drug. We visualised the results in Fig. 3a and Supp. Fig. 4a (PA). The results were corroborated with existing literature. Importantly, GeneBac has no prior information on the resistant genes for each drug, so it has to learn it on its own during training. We believe that finding genes associated with resistance is a crucial step to understanding the molecular mechanisms of antibiotic resistance, which could be also validated with for example a knock-out experiment.

##### Leveraging GNN Explainer for Graph Neural Network Interpretability

To provide interpretable explanations for the GNN integrating interactions between genes, we used GNN Explainer [41], which given an input example identifies a compact subgraph structure and a small subset of node features that have a crucial role in GNN’s predictions. Here a node represents a gene with its node features computed using the CNN with the DNA sequence of the gene as input, while the edges of the graph were extracted from the STRING DB based on the prior knowledge of interactions between genes. We used GeneBac trained on the MIC prediction task using the MTB CRyPTIC dataset. The node features were computed using the trained CNN, which were then fed into the GNN Explainer. We used the implementation provided as part of the PyTorch Geometric [59] package and set the number of epochs in the GNN Explainer to 200 and the learning rate to 0.01 which is the default. We masked entire nodes and edges, rather than specific features (”object” mask type value) and set the explanation type to explain the model predictions. Finally, we set the task in the model config to the graph level.

We ran the GNN Explainer on the held-out strains from the MTB CRyPTIC dataset and collected the node mask as well as the edge mask. We used the node mask

to visualise the aggregate normalised mean node weights across samples (Supp. Fig. 5). We also visualised the subgraph structure for several samples (Supp. Fig. 5c-f). In these visualisations, the size of the node indicates the node weight and the edges with high weight are highlighted.

##### Gene expression prediction

We used the dataset of 386 strains with paired DNA and RNA-seq reads [42] for the genes in the *Pseudomonas aeruginosa* core genome (4,548 genes). We treated each gene in each strain as a separate example, allowing the model to learn common DNA features across different genes. We randomly split the dataset into train (70%), validation (10%) and test (20%) across strains.

The CNN used for gene expression prediction was the same as for the antibiotic resistance prediction model, with the addition of the feedforward layer which takes in as input a gene representation and projects it into a scalar value, used as the gene expression prediction.

##### Categorising gene variance

We computed the gene variance based on the variation in the gene expression across strains. We computed the standard deviation across strains for each gene and based on the distribution of the standard deviation, we have set the following thresholds: (1) high variance genes are genes whose standard variation is 0.7, (2) medium variance genes have standard deviation between 0.7 and 0.4 and (3) low variance genes which have standard deviation below 0.4. The majority of genes belong to the low-variance gene class.

##### Benchmarking gene expression prediction

As a benchmark for GeneBac, we used a mean of the gene expression in a given lineage. We did it by aggregating the strains across lineages and genes using the training set. Finally, we computed the mean expression of each gene and used it for prediction during evaluation.

##### Motif discovery

To examine whether the gene expression model recovers TF binding motifs, we used DeepLift [34] to compute importance scores per nucleotide using the PA reference genome PAO1. We then leveraged TF-MoDISco [43] and a set of position-weight matrices (PWM) of TF binding motifs in PA to match the motifs extracted from the model’s importance scores with the database of PA motifs. We discarded the patterns matched by TF-MoDISco which had the *q-value* higher than 0.05 to remove the low confidence matches. Matched motifs together with number of motif instances found on the genome are visualised on the Fig. 4e. We also visualised the distribution of number of motif binding sites per gene 4f, excluding the genes with no predicted motifs.

The PWM database was established by combining PA motifs from Prodoric [60] and CollecTF databases [61]. Importantly, TF-MoDISco relies on high-quality PWMs

and we have observed that the overlap between the motifs in different databases can vary significantly, pointing out the limitation of such studies. We share the set of PWMs used in the study as part of the GitHub repository.

##### Computing relative base-pair importance

Certain base pair positions across genes tend to particularly affect the RNA abundance of the gene, an example of such is the promoter, or the start codon, around which a mutation may lead to a change in the expression of a gene. To evaluate whether the base-pair importance is enriched in particular regions of the gene, we computed the absolute base pair importance for the core genes in the reference genome PAO1 using DeepLift [34]. We then normalised the base-pair importance by the gene length, as genes have strongly varying lengths. Using the mean importance at each base pair position, we visualised the result in the Fig. 4g.

##### Gene representation visualisation

To examine whether the genes cluster by their position on the chromosome, we extracted the low dimensional (64) gene representations from the validation set of the gene expression dataset. To get a single embedding for each gene, we took the mean of all embeddings of the same gene. We used the start codon of a gene from the PAO1 reference genome as the position on the chromosome. We used t-SNE [33] to visualise the embeddings in two dimensions. We set the perplexity of the t-SNE algorithm to 20 and the number of iterations to 1500.

---

**Supp. Fig. 1.**
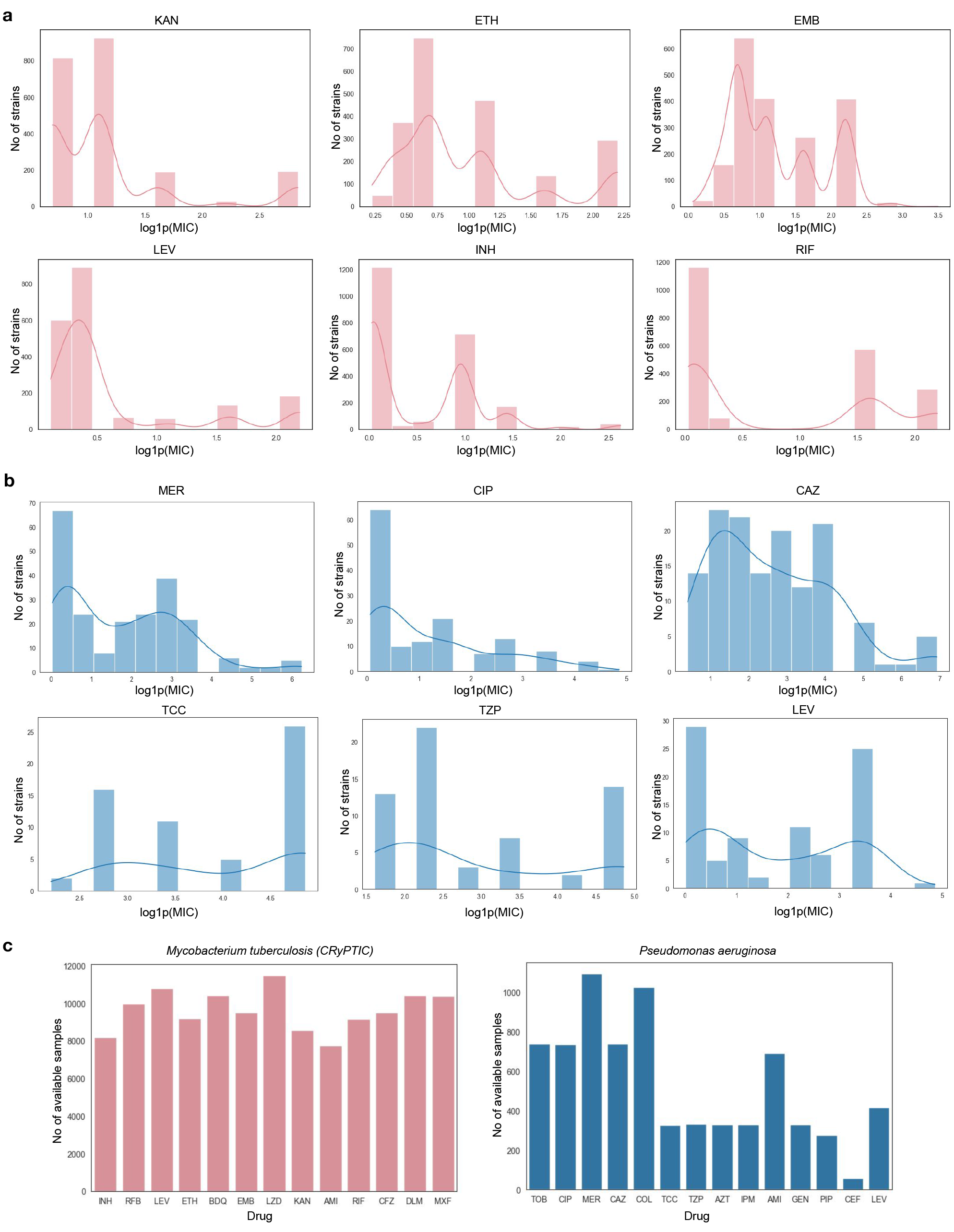
Dataset statistics. *(a)* MIC distribution for a set of drugs from the *Mycobacterium tuberculosis* (CRyPTIC) dataset. *(b)* MIC distribution for a set of drugs from the *Pseudomonas aeruginosa* dataset. *(c)* Number of samples available per drug on the *Mycobacterium tuberculosis* and *Pseudomonas aeruginosa* datasets.

**Supp. Fig. 2.**
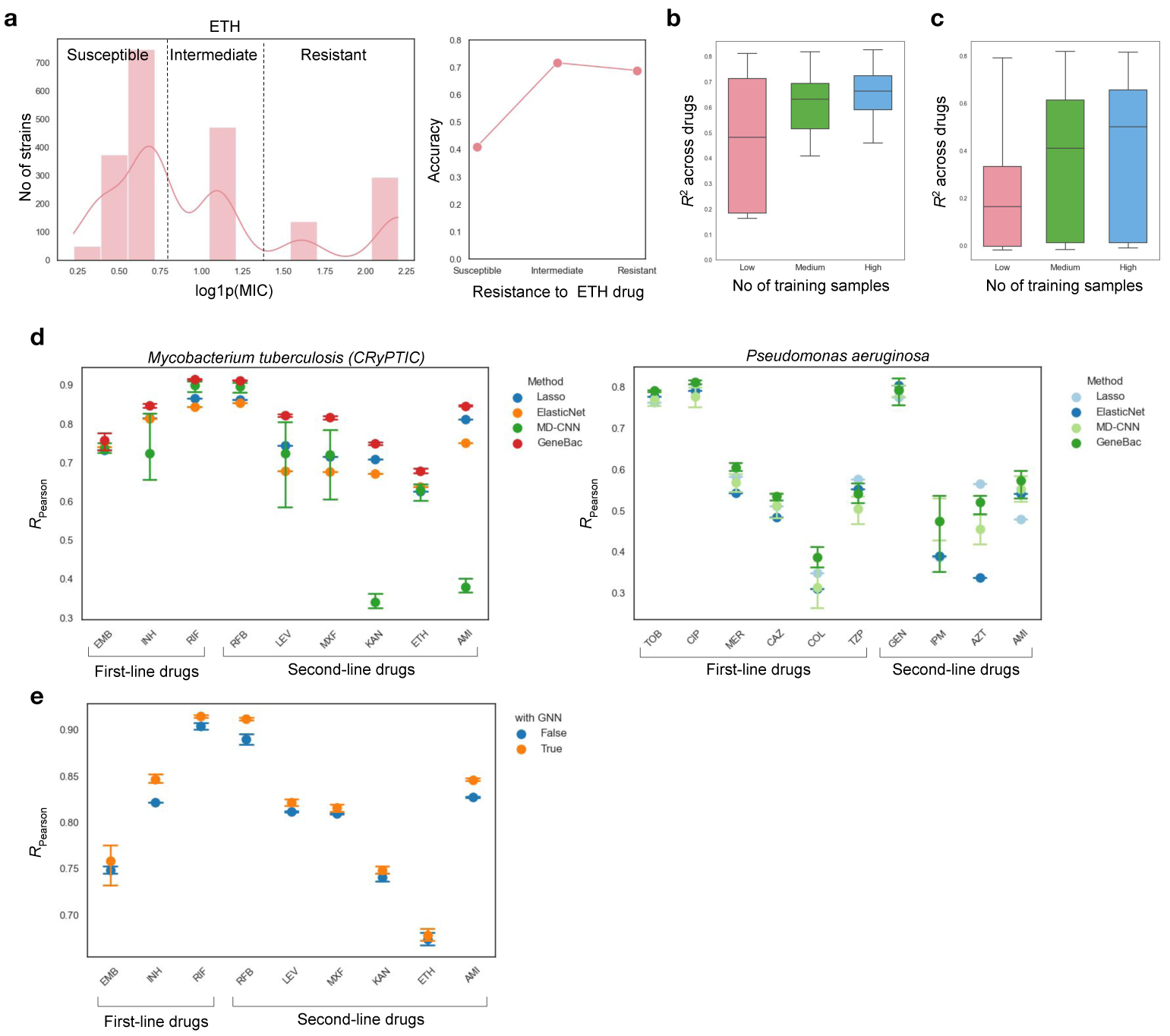
Additional performance comparison on the antibiotic resistance MIC prediction task. *(a)* The observed MIC distribution of the Ethionamide drug on the MTB dataset divided into 3 categories: susceptible, intermediate and resistant (left). Accuracy for each category on held-out strains of the GeneBac model trained on the MIC prediction task (right). *(b)* Performance across drugs on different training data regimes on the MTB dataset across all drugs. Here, low, medium and high number of samples means using 25%, 50% and 100% training samples available. Each box represents the performance across drugs, with the horizontal line representing median performance. *(c)* Performance across drugs on different training data regimes on the MTB dataset across non- first-line drugs. Here, low, medium and high number of samples means using 25%, 50% and 100% training samples available. Each box represents the performance across drugs, with the horizontal line representing median performance. *(d)* Performance comparison between different methods on first- and second-line drugs on the MIC prediction regression task. The error bars represent the variance across runs and the dot represents mean performance. *(e)* The impact of the GNN on performance across drugs compared to a baseline method which uses a dense layer on top of mean pooled gene representations on the MTB dataset. The error bars represent the variance across runs and the dot represents mean performance.

**Supp. Fig. 3.**
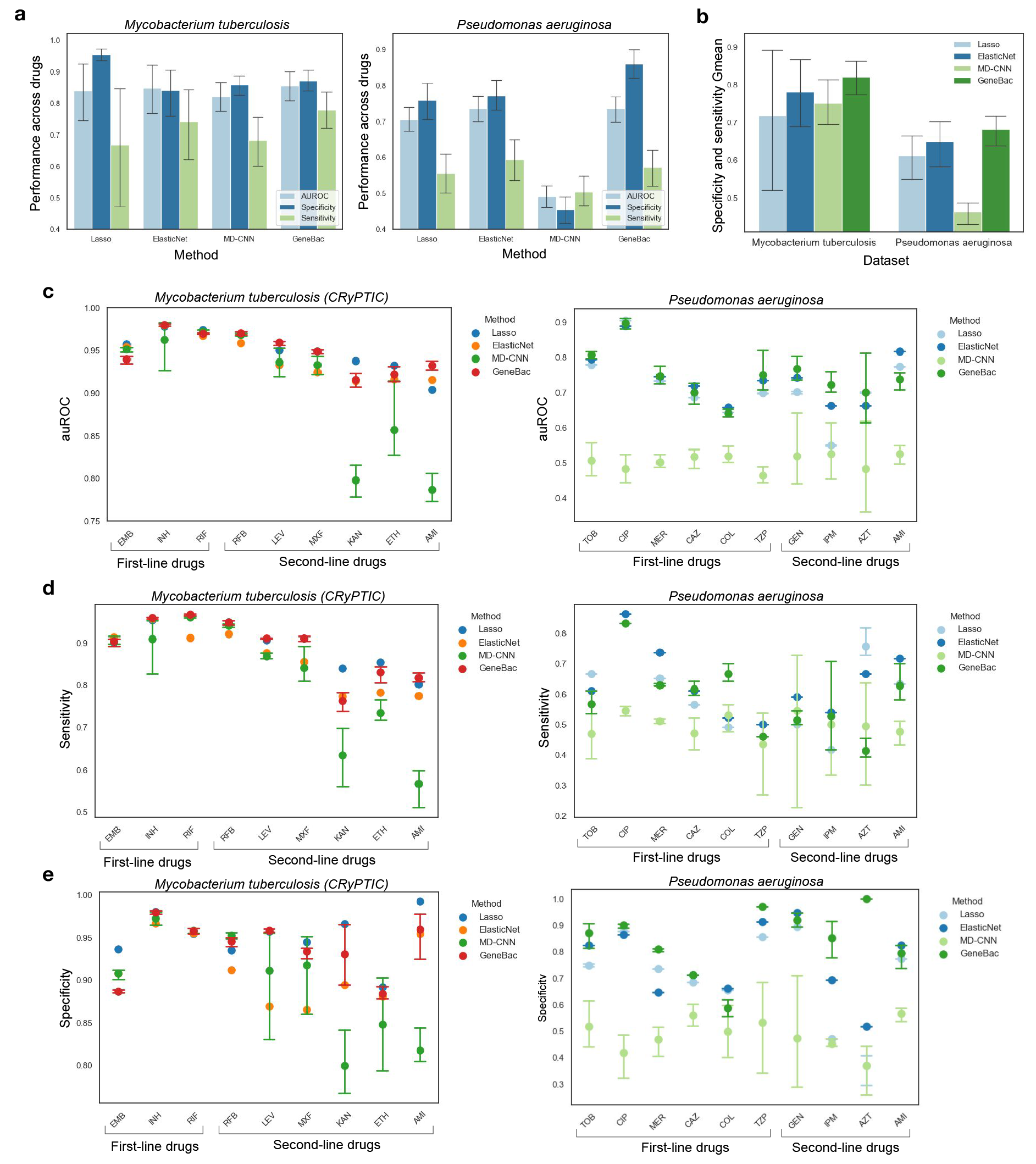
Performance on the binary antibiotic resistance prediction task. *(a)* Performance com- parison of GeneBac and other methods on the antibiotic resistance binary prediction task evaluated by binary classification metrics on all drugs. The error bars represent the variance across runs and the color bars represent the mean performance across different metrics.*(b)* Performance comparison of GeneBac and other methods on the antibiotic resistance binary prediction task evaluated by the geometric mean of specificity and sensitivity on all drugs and two different datasets. The error bars represent the variance across runs and the color bars represent the mean performance. *(c)* Perfor- mance comparison between different methods on first- and second-line drugs on the binary antibiotic resistance prediction task as evaluated by the area under the ROC curve (AUROC). The error bars represent the variance across runs and the dot represents mean performance. *(d)* Performance com- parison between different methods on first- and second-line drugs on the binary antibiotic resistance prediction task as evaluated by Sensitivity. The error bars represent the variance across runs and the dot represents mean performance. *(e)* Performance comparison between different methods on first- and second-line drugs on the binary antibiotic resistance prediction task as evaluated by specificity. The error bars represent the variance across runs and the dot represents mean performance.

**Supp. Fig. 4.**
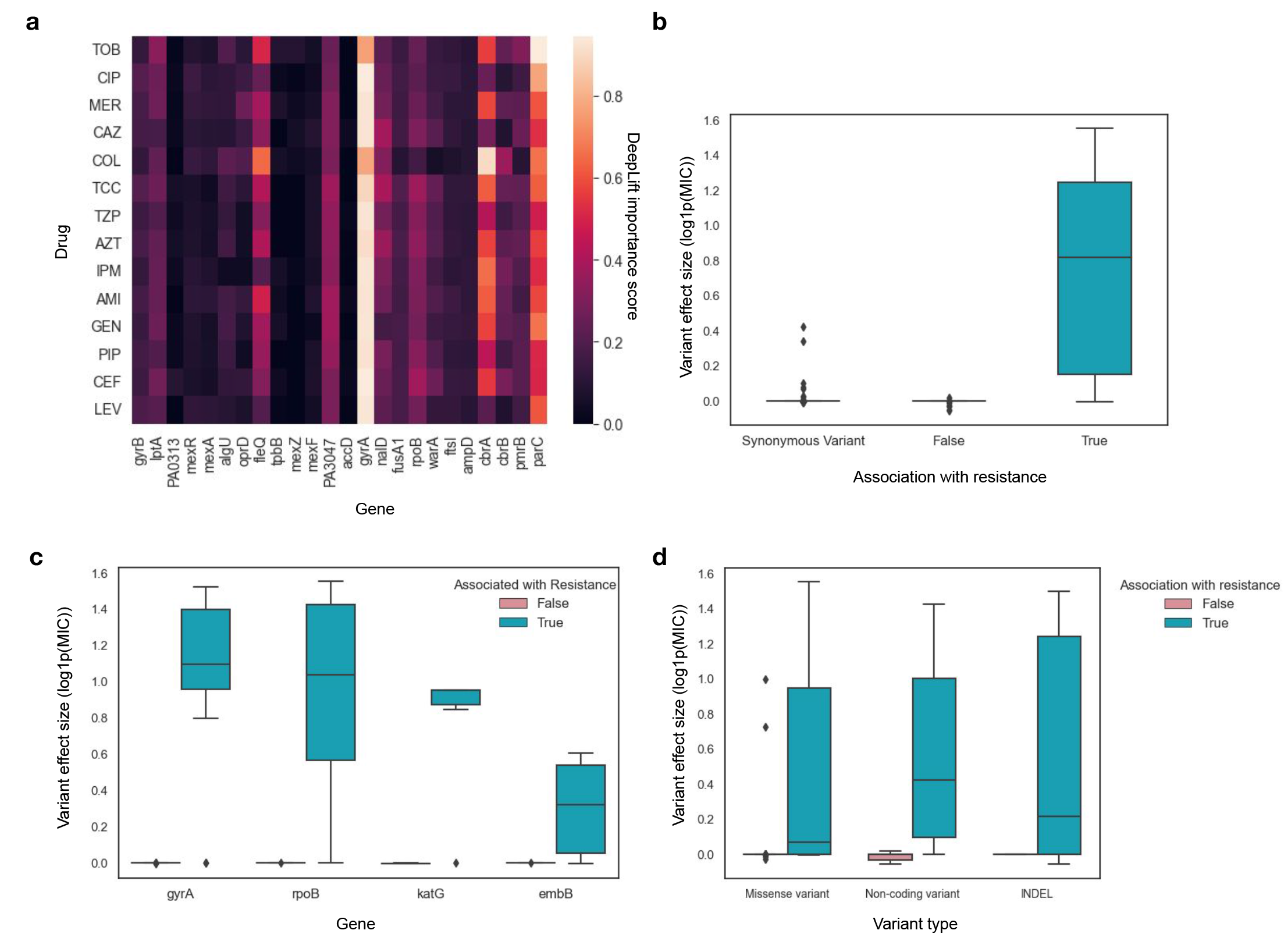
Additional GeneBac variant effect prediction results. *(a)* Drug-Gene importance mea- sured by DeepLift importance score, aggregated across held-out strains and normalised per target (i.e. gene) on the PA dataset. *(b)* Variant effect size of variants from the WHO MTB mutation cata- logue on variants associated with resistance, not associated with resistance and synonymous variants. The variant effect size was calculated using the GeneBac model trained on the MIC regression pre- diction task on the MTB dataset. Each box represents the variant effect size across different variants with the horizontal line representing the median variant effect size. *(c)* Variant effect size of vari- ants from the WHO MTB mutation catalogue across a set of genes on variants associated and not associated with resistance. Each box represents the variant effect size across different variants with the horizontal line representing the median variant effect size. *(d)*Variant effect size of variants from the WHO MTB mutation catalogue across different variant types. To allow for meaningful analysis across different mutation types, we included low-confidence variants from the catalogue marked as *interim*. Each box represents the variant effect size across different variants with the horizontal line representing the median variant effect size.

**Supp. Fig. 5.**
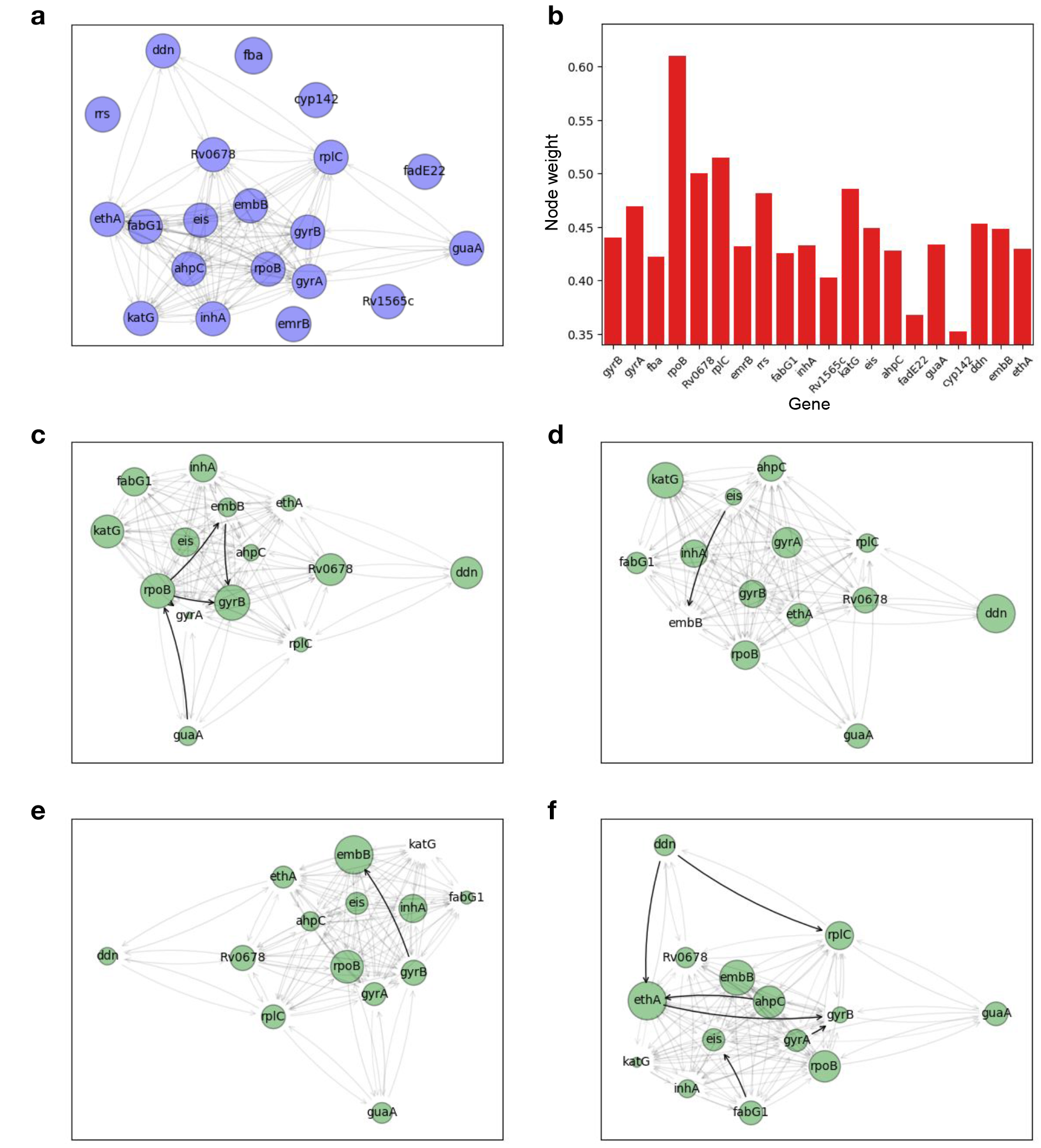
Graph Neural Network models interactions between genes. *(a)* The graph repre- senting interactions (edges) between selected genes (nodes) in *Mycobacterium Tuberculosis* used for training and evaluation of the model. *(b)* Node weights computed using the GNN Explainer on the *Mycobacterium Tuberculosis* CRyPTIC dataset on held-out strains. High node weight indicates high importance. *(c)* GNN Explainer indicates rpoB as a key gene for final prediction based on edge and node weights in a *Mycobacterium Tuberculosis* strain *site.01.subj.DR0013.lab.DR0013.iso.1* from the CRyPTIC dataset with high resistance to Rifampicin. *(d)* GNN Explainer indicates katG as a key gene for final prediction based on node weights in a *Mycobacterium Tuberculosis* strain *site.01.subj.DR0114.lab.DR0114.iso.1* from the CRyPTIC dataset with high resistance to Kanamycin. *(e)* GNN Explainer indicates embB as a key gene for final prediction based on edge and node weights in a *Mycobacterium Tuberculosis* strain *site.01.subj.DR0163.lab.DR0163.iso.1* from the CRyPTIC dataset with high resistance to Ethambutol. *(f)* GNN Explainer indicates ethA as a key gene for final prediction based on edge and node weights in a *Mycobacterium Tuberculo- sis* strain *site.02.subj.0353.lab.235092-15.iso.1* from the CRyPTIC dataset with high resistance to Ethionamide.

**Supp. Fig. 6.**
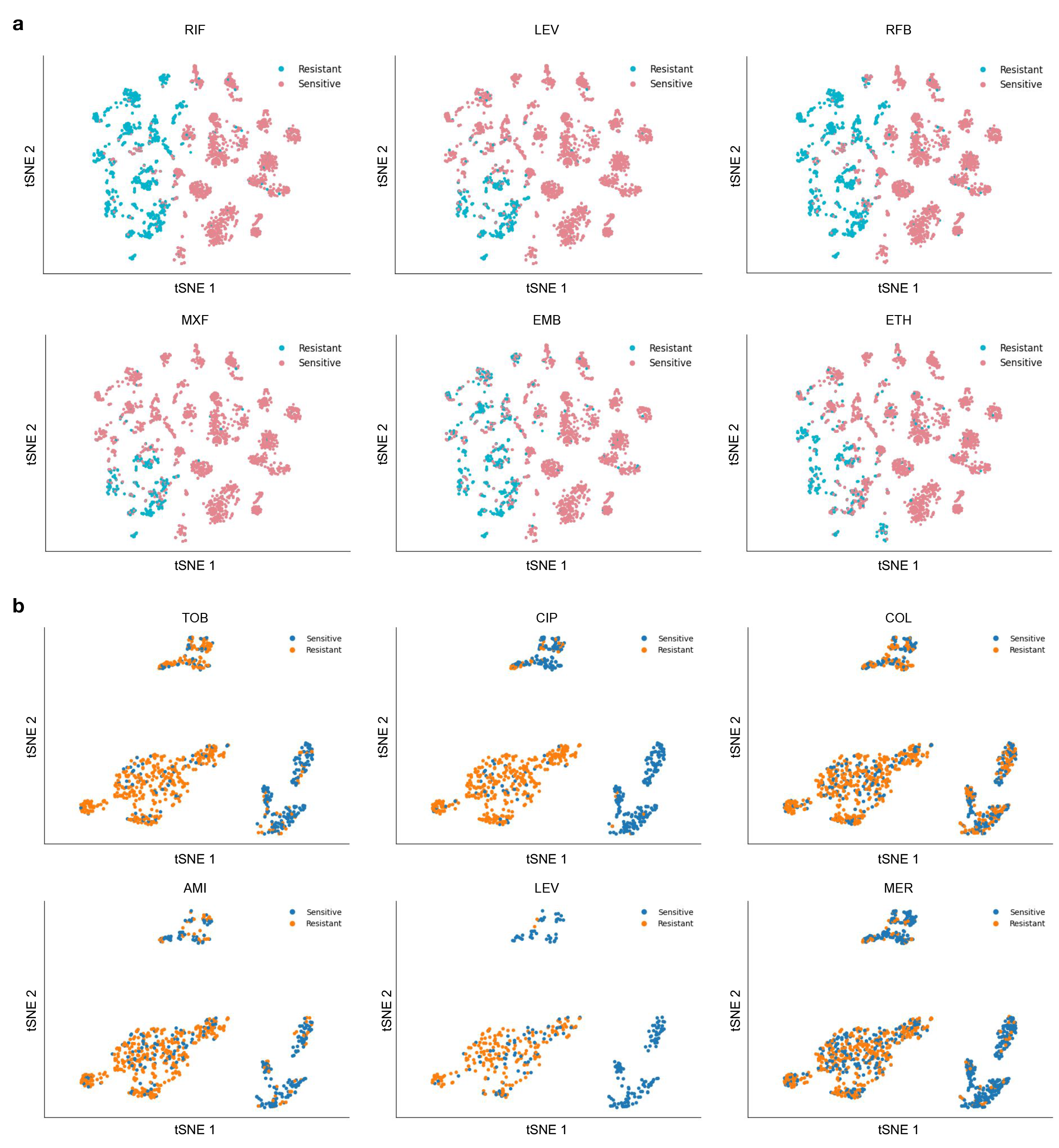
Additional strain representation clustering visualisations. *(a)* Strain embeddings learned on the MTB dataset coloured by their resistance to a drug. *(b)* Strain embeddings learned on the PA dataset coloured by their resistance to a drug.

**Supp. Table 1.** *Mycobacterium tuberculosis* genes selected as model input including their location on the H37Rv reference genome and association with drugs available in the dataset.

| Gene | Start | End | Drug(s) | Length |
| --- | --- | --- | --- | --- |
| gyrB | 5140 | 7267 | Ethambutol, Levofloxacin | 2118 |
| gyrA | 7202 | 9818 | Ethambutol, Amikacin, Ethionamide, Kanamycin, Levofloxacin, Moxifloxacin | 2616 |
| fba | 441265 | 442399 | Delamanid | 1134 |
| rpoB | 759707 | 763325 | Ethambutol, Isoniazid, Rifampicin, Amikacin, Ethionamide, Kanamycin, Levofloxacin, Moxifloxacin, Rifabutin, Bedaquiline, Linezolid | 3618 |
| Rv0678 | 778890 | 779487 | Bedaquiline, Clofazimine | 597 |
| rplC | 800709 | 801462 | Linezolid | 753 |
| emrB | 876818 | 878540 | Linezolid | 1722 |
| rrs | 1471746 | 1473382 | Amikacin, Kanamycin, Levofloxacin, Moxifloxacin, Bedaquiline | 1636 |
| fabG1 | 1673340 | 1674183 | Isoniazid, Ethionamide, Clofazimine | 843 |
| inhA | 1674102 | 1675011 | Isoniazid, Ethionamide | 909 |
| Rv1565c | 1771640 | 1773929 | Rifampicin | 2289 |
| katG | 2153889 | 2156211 | Ethambutol, Isoniazid, Rifampicin, Moxifloxacin, Rifabutin | 2322 |
| eis | 2714124 | 2715432 | Kanamycin | 1308 |
| ahpC | 2726093 | 2726780 | Isoniazid | 687 |
| fadE22 | 3423262 | 3425527 | Delamanid | 2265 |
| guaA | 3812501 | 3814178 | Rifampicin, Amikacin | 1677 |
| cyp142 | 3954325 | 3955621 | Clofazimine | 1296 |
| ddn | 3986744 | 3987299 | Delamanid | 555 |
| embB | 4246414 | 4249810 | Rifampicin, Levofloxacin, Moxifloxacin, Rifabutin | 3396 |
| ethA | 4326004 | 4327373 | Ethionamide, Kanamycin | 1369 |

**Supp. Table 2.** Hyperparameters used for the antibiotic resistance prediction task (both regression and binary). Where two values are available, the parameter in bold indicates the parameter used on the MTB dataset and the underlined one on the PA dataset.

|  |  |
| --- | --- |
| Number of CNN layers | 7 |
| Number of GNN layers | 2 |
| Gene representation embedding | 64 |
| Strain representation embedding | 64 |
| Max gene length | 2560 |
| Max region upstream of the gene length | **100**, <u>200</u> |
| Learning rate | 0.0001, 0.0005 |
| Dropout | 0.2 |
| Optimiser | AdamW |
| Optimiser momentum | $\beta_1 = 0.9$, $\beta_2 = 0.999$ |
| LR scheduler | Linear with warmup |
| % of warmup steps | 0.1 |
| Batch size | 16 |
| Reverse complement aug. % | 0.5 |
| Gradient clipping value | 1.0 |
| Max epochs | 500 |

**Supp. Table 3.** Hyperparameters used for the gene expression task.

|  |  |
| --- | --- |
| Number of CNN layers | 7 |
| Gene representation embedding | 64 |
| Max gene length | 2560 |
| Max region upstream of the gene length | 200 |
| Learning rate | 0.0001 |
| Dropout | 0.2 |
| Optimiser | AdamW |
| Optimiser momentum | $\beta_1 = 0.9$, $\beta_2 = 0.999$ |
| LR scheduler | Linear with warmup |
| % of warmup steps | 0.1 |
| Batch size | 256 |
| Reverse complement aug. % | 0.5 |
| Gradient clipping value | 1.0 |
| Max epochs | 100 |

**Supp. Table 4.** Details on drugs used in the study.

| *Mycobacterium tuberculosis* |  |  | *Pseudomonas aeruginosa* |  |  |
| --- | --- | --- | --- | --- | --- |
| Drug | Drug code | Drug line | Drug | Drug code | Drug line |
| Moxifloxacin | MXF | Second | Tobramycin | TOB | First |
| Bedaquiline | BDQ | New and repurposed | Ciprofloxacin | CIP | First |
| Kanamycin | KAN | Second | Meropenem | MER | First |
| Clofazimine | CFZ | New and repurposed | Ceftazidime | CAZ | First |
| Amikacin | AMI | Second | Colistin | COL | First |
| Delamanid | DLM | New and repurposed | TicarClav | TCC | New and repurposed |
| Rifabutin | RFB | Second | PiperacillinTazobactam | TZP | First |
| Linezolid | LZD | New and repurposed | Aztreonam | AZT | Second |
| Ethambutol | EMB | First | Imipenem | IPM | Second |
| Levofloxacin | LEV | Second | Amikacin | AMI | Second |
| Ethionamide | ETH | Second | Gentamicin | GEN | Second |
| Isoniazid | INH | First | Piperacillin | PIP | New and repurposed |
| Rifampicin | RIF | First | Cefepime | CEF | New and repurposed |
|  |  |  | Levofloxacin | LEV | New and repurposed |

